# Laboratory and wild *Drosophila sechellia* have conserved niche specialization phenotypes

**DOI:** 10.64898/2026.01.26.701819

**Authors:** Michael P. Shahandeh, Liliane Abuin, Olakunle A. Jaiyesimi, Polpass Arul Jose, Suhrid Ghosh, Asfa Sabrin Borbora, Janpreet Kaur, Cassandra G. Extavour, Richard Benton

## Abstract

A major challenge to investigating the proximate causes of ecological adaptation is the difficulty of studying the phenotypes of organisms in their natural environments. By necessity, many studies seeking to determine the genetic and cellular basis of adaptation therefore investigate potentially adaptive phenotypes under laboratory conditions where organisms are more easily experimentally manipulated. For laboratory models, it remains unclear if organisms maintained long term under laboratory conditions are representative of relatives in their natural environment. In recent years, *Drosophila sechellia*, a specialist species endemic to the Seychelles, has emerged as a (neuro)genetic model for studying the molecular basis of ecological adaptation. A multitude of studies have investigated the genetic and cellular basis of various aspects of this species’ specialization in a laboratory setting. However, the vast majority of these studies use laboratory strains of *D. sechellia* that were collected many decades ago, and have been maintained under conditions very different from their natural niche. Thus, it remains unclear if and how these strains resemble their wild counterparts. Here, we compare the phenotypes of these laboratory strains with recently-collected wild *D. sechellia* to ask if laboratory strains display a loss or degradation of phenotypes potentially involved in their specialization resulting from their long-term laboratory maintenance. Across several behavioral and anatomical phenotypes, we find a high degree of similarity between wild-caught and laboratory-maintained strains. Our results suggest that studies of the molecular mechanisms underlying *D. sechellia*’s phenotypes associated with specialization are likely representative of the evolution of these flies in the wild.

## Introduction

A significant challenge to studying the genetic and cellular underpinnings of ecological adaptation is the necessity to remove individuals from their natural niche for laboratory experimentation. This means that many mechanistic studies of the development, physiology and behavior of a wide variety of organisms are traditionally performed in conditions that differ, sometimes drastically, from an animal’s natural environment. This issue is particularly pertinent for laboratory model organisms, where strains are maintained for generations in artificial environments, creating the potential for adaptation to laboratory conditions, which results in uncertainty as to whether phenotypes measured in the laboratory are representative of those in nature. For instance, a standard *Caenorhabditis elegans* laboratory strain displays dramatic differences in social and feeding behaviors, growth, and reproductive output compared to wild *C. elegans*, likely resulting from adaptation to laboratory rearing conditions^1,2^. Moreover, several laboratory strains of *C. elegans,* and even other *Caenorhabditis* species, have convergently evolved unnatural resistance to dauer-inducing pheromones – a starvation-resistant stage of developmental arrest – which may be an adaptation to laboratory culture at high densities^3^. Exacerbating this problem, it is possible for the same genetic isolates that are maintained for years in different laboratories to experience different selection pressures resulting from differences between these environments (e.g., culture medium and rearing practices) creating laboratory-specific phenotypic differences for the same original laboratory strain. For example, laboratory-specific replicates of the commonly used *Drosophila melanogaster* isogenic strain, Canton-S, display significant differences in locomotor behavior^4^. Likewise, in the mouse model, *Mus musculus*, laboratory rearing environment influences phenotypic measures across biological levels^5^, in some cases resulting in the division of genetic isolates into substrains with known phenotypic differences^6,7^.

The issue of phenotypic divergence upon prolonged laboratory rearing is potentially more extreme for specialist species, which display adaptation to a narrow ecological niche that is not always easily replicated in a laboratory setting. One example is *Drosophila sechellia*, which diverged from a common ancestor with *D. melanogaster* no more than 5 million years ago^8^. Despite the phylogenetic proximity and high genomic and anatomical similarity of these drosophilids^9^, *D. sechellia* occupies a dramatically different ecological niche. While *D. melanogaster* is a cosmopolitan species, feeding generally on rotting fruit^10^, *D. sechellia* is endemic to the Seychelles archipelago, where it specializes on the ripe noni fruit of the shrub *Morinda citrifolia*^11^ (Figure 1A), which is toxic to most other *Drosophila* species^12^. Moreover, in its near-equatorial habitat, *D. sechellia* is exposed to tropical conditions year-round, unlike globally-distributed *D. melanogaster*^8^, which experiences higher degrees of seasonal variation at high latitudes. As such, *D. sechellia* has been a focus of many comparative studies aimed at describing the genomic, developmental, physiological, behavioral, and neurological evolution that underlies its niche specialization (reviewed in^13^).

**Figure 1.**
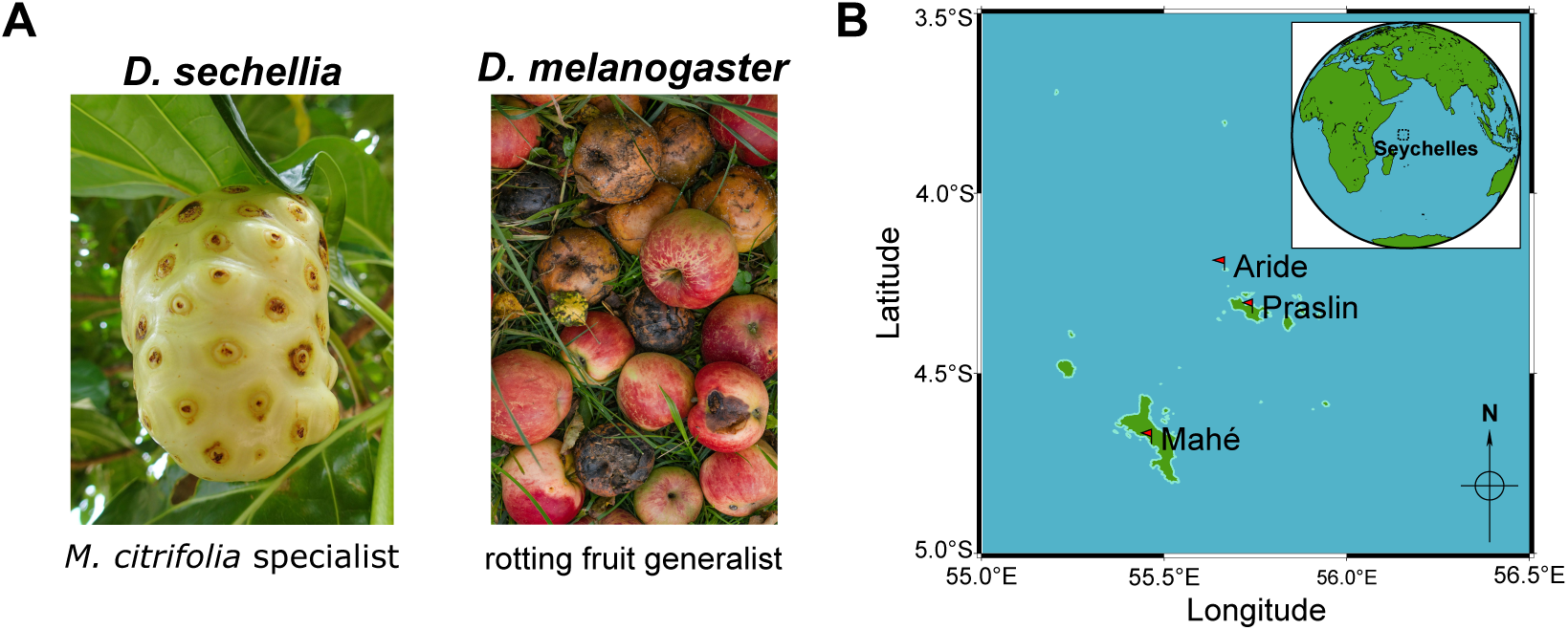
*D. sechellia* is a *M. citrifolia* specialist endemic to an equatorial island archipelago. **A.** Host preferences of *D. melanogaster* and *D. sechellia*. **B.** A map of the Seychelles archipelago with specific coordinates of wild *D. sechellia* strain collection sites on three islands marked with red flags (Mahé Island coordinates: 4.69424° S, 55.46580° E, Aride Island coordinates: 4.21423° S, 55.66555° E, and Praslin Island coordinates: 4.33340° S, 55.74180° E). Upper-right inset shows the location of the Seychelles on the eastern hemisphere. The maps were generated using Generic Mapping Tools (GMT) version 6.6.0 (Wessel et al 2019).

Most studies on *D. sechellia* have used laboratory adapted strains collected decades ago (with approximately 25-30 generations per year). Additionally, in the laboratory, these strains typically are not reared on their specific host fruit, but rather on standard *Drosophila* medium sometimes supplemented with commercially available noni juice. This juice only partially resembles noni fruit chemically^14^, and has limited, at most, toxicity for other drosophilids (unpublished observations). It is unclear how representative these laboratory strains of *D. sechellia* are of those living in their specific niche today, or whether specialization has degraded over the course of laboratory maintenance and adaptation. Recent field studies have sought to address similar issues in *D. melanogaster* by observing the behaviors of natural populations of flies to collect data that can be compared with laboratory generated data^15–17^. Here, we also sought answers to these questions by performing developmental, physiological, and behavioral laboratory assays on recently collected lines of *D. sechellia* from different islands of the Seychelles archipelago (Figure 1B). We measured multiple aspects of *D. sechellia*’s host specialization to compare the behavior of these non-laboratory adapted and laboratory strains.

## Results

### Wild *D. sechellia* collection

*D. sechellia* flies were collected in April 2023 during a field trip to three different islands in the Seychelles archipelago (Figure 1B) and used to found isofemale lines. Strain M1311 was obtained from the grounds of the National Biodiversity Center on Mahé Island. Strains A3312 and A3F311 were collected from a site on Aride Island. Strains P7611 and P7621 were collected from outside Vallee de Mai on Praslin Island. All strains except A3F311 were founded from single adult females collected directly from the surfaces of fallen noni fruits using a manual fly aspirator. Strain A3F311 was founded from a female that eclosed from a larval-infested fallen noni fruit collected in the field and brought back to the laboratory. Each strain was determined to be *D. sechellia* by molecular barcoding (see Methods).

### Olfactory preference for ripe noni volatiles

*D. sechellia* displays a strong preference for ripe noni fruit and noni-derived acids over other substrates in open trap assays permitting olfactory and gustatory sampling both in the lab^18^ and field^19^. In long-range and short-range laboratory assays that exclude gustatory sampling, *D. sechellia* displays increased olfactory-mediated preference for noni and noni volatiles^14,20,21^. This strong preference for noni has been attributed, in part, to evolution of the peripheral olfactory sensory system^20,22^.

To compare the phenotypes of laboratory and wild *D. sechellia*, we first examined their olfactory preference given a choice between noni fruit juice and apple cider vinegar (a fermentation product that is highly attractive to *D. melanogaster*^23^). We tested two commonly-used laboratory *D. sechellia* strains, the five new wild-caught *D. sechellia* strains and one *D. melanogaster* laboratory strain. In our trap design, flies could not physically encounter the liquid prior to making a choice, limiting the possibility for gustatory sensation (Figure 2A). All seven *D. sechellia* strains, both laboratory and recently-caught, displayed significant preference for noni juice over apple cider vinegar, while *D. melanogaster* displayed a slight but non-significant preference for apple cider vinegar (Figure 2B). Importantly, there were no distinguishable differences when comparing the preferences between *D. sechellia* strains, regardless of their collection date. However, all *D. sechellia* strains displayed significantly different preference indices from *D. melanogaster*.

**Figure 2.**
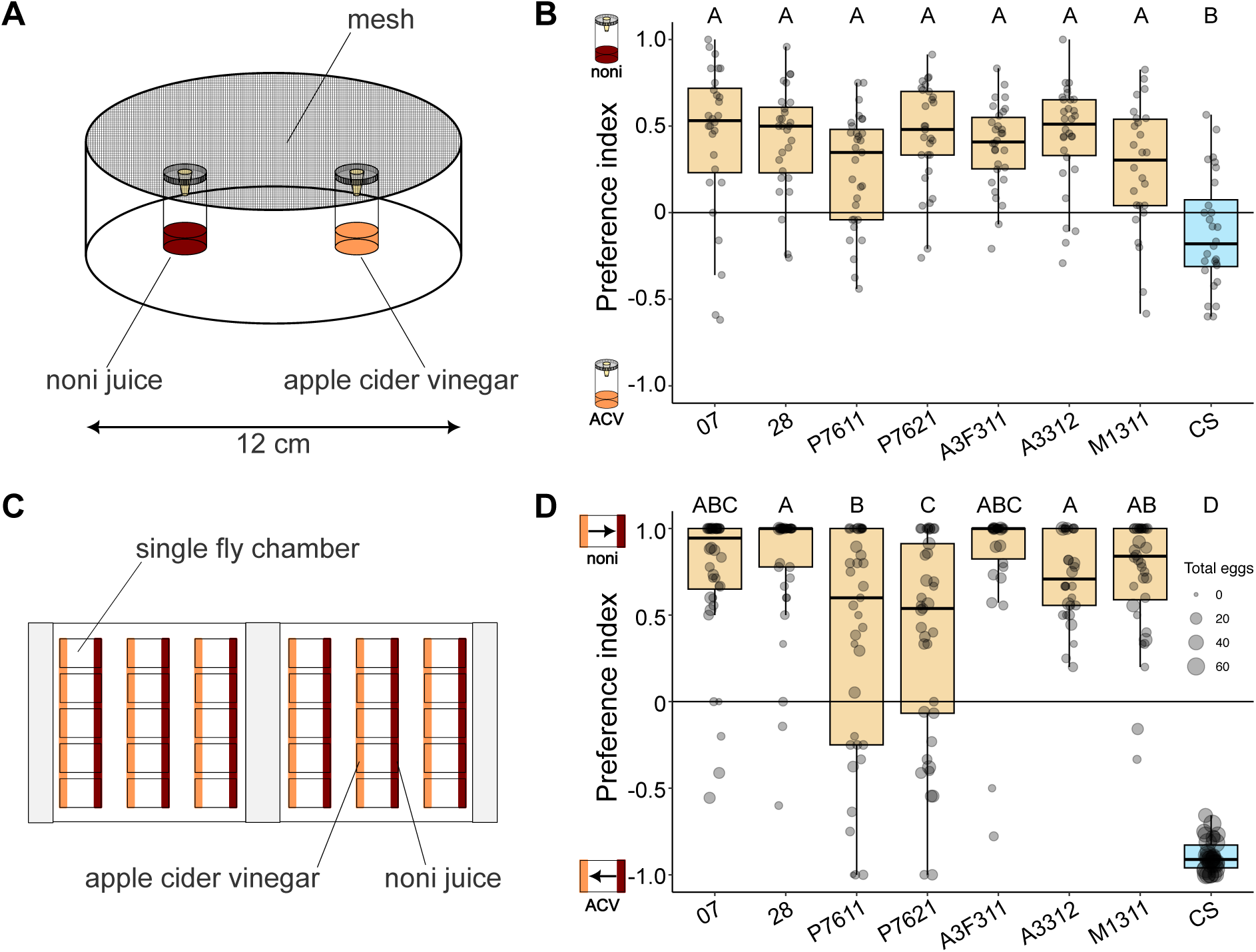
*D. sechellia* display strong olfactory preference for noni in short-range olfactory and oviposition assays. **A.** Schematic of the short-range olfactory preference assay. **B.** Short-range olfactory preference of wild and laboratory *D. sechellia* strains and *D. melanogaster* Canton-S (CS). All *D. sechellia* strains displayed a significant preference for noni juice over apple cider vinegar (all p < 0.01, Wilcoxon test), while CS displayed no significant preference. Sample sizes: 07 (28), 28 (28), P7611 (31), P7621 (29), A3F311 (30), A3312 (30), M1311 (26), and CS (28). **C.** Schematic of the single-fly oviposition preference assay. **D.** Single-fly oviposition preference for the same strains. Point size is scaled by the total number of eggs laid per female. All *D. sechellia* strains displayed a significant preference for noni juice over apple cider vinegar (all p < 0.01, Wilcoxon test), while CS displayed a significant preference for apple cider vinegar (p < 0.001, Wilcoxon test). Sample sizes: 07 (40), 28 (39), P7611 (37), P7621 (37), A3F311 (30), A3312 (30), M1311 (36), and CS (34). For panels **B** and **D**, letters above boxplots indicate pairwise significant differences (all p < 0.01, Wilcoxon test followed by Bonferroni correction).

### Oviposition preference for noni substrates

Upon successful location of a host fruit, flies will feed, engage in social behaviors with other flies they encounter and reproduce^15^. *D. sechellia* consistently selects ripe noni or noni media for oviposition over other substrates^18,19,24,25^. This inherent preference to oviposit on ripe noni has been attributed to olfactory receptor tuning changes^24^ and gustatory preference differences^26^.

To compare the oviposition preferences of our recently-caught and laboratory *D. sechellia* strains, we replicated a single-fly oviposition choice assay^27^ ^24^. Briefly, we offered individual gravid females a choice between two oviposition media of equivalent texture, made with noni juice or apple cider vinegar, over a 24-hour period. During this period, flies were able to sample each media using tactile, gustatory, and olfactory cues (Figure 2C). In this assay, all *D. sechellia* strains displayed a significant preference for ovipositing on noni media over apple cider vinegar media, while our *D. melanogaster* laboratory strain displayed a significant preference for ovipositing on apple cider vinegar media (Figure 2D). All *D. sechellia* strains were statistically distinguishable from *D. melanogaster*. In this assay we observed significant variation among our *D. sechellia* strains with respect to the strength of their preference for ovipositing in noni media, but this did not reflect a consistent distinction between laboratory and wild *D. sechellia* (Figure 2D).

### Olfactory sensory neuron population expansion and contraction

The species-specific host location and oviposition behaviors of *D. sechellia* are thought to be due in part to increases in the numbers of olfactory sensory neurons (OSNs) detecting characteristic noni volatiles^28^, namely hexanoic acid-sensing Ir75b OSNs in antennal coeloconic 3I (ac3I) sensilla, and Or22a and Or85b OSNs in antennal basiconic 3 (ab3) sensilla, which detect methyl esters and *2*-heptanone, respectively^14,29^ ^20^.

To determine whether the expansions described in laboratory strains are a conserved phenotype of *D. sechellia* in the wild, we used immunofluorescence or RNA fluorescence *in-situ* hybridization to label specific OSN populations in one laboratory strain each of *D. melanogaster* and *D. sechellia*, as well as three of the wild-caught *D. sechellia* strains (P7621, A3312, and M1311). Across both laboratory and wild-caught *D. sechellia* strains, we observed a consistent two-fold increase in Ir75b OSNs relative to *D. melanogaster* (Figure 3A). Similarly, by labelling Or85b OSNs, we detected a consistent expansion of ab3 sensilla (Figure 3B), with *D. sechellia* strains exhibiting a three-fold increase in OSN number compared to *D. melanogaster*.

**Figure 3.**
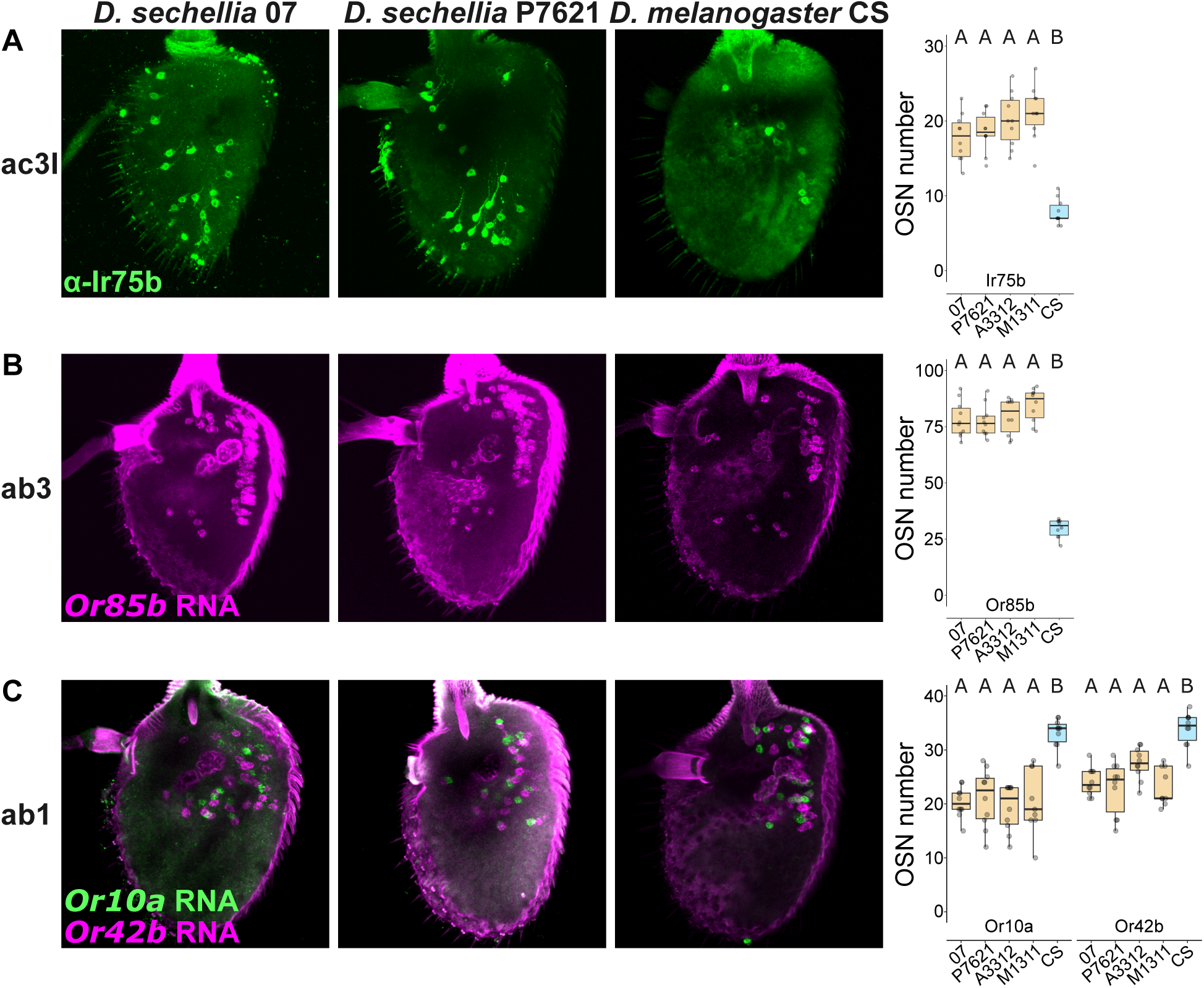
Olfactory sensory neuron population expansion and contraction in *D. sechellia*. **A.** Left: example images of Ir75b OSNs in ac3I sensilla of the third antennal segment of laboratory *D. sechellia* (07), wild *D. sechellia* (P7612), and *D. melanogaster* (CS). Right: quantifications of Ir75b OSNs in laboratory *D. sechellia* (07), *D. melanogaster* (CS), and three wild *D. sechellia strains*, one from each island (P7612, A3312, M1311). **B.** Left: example images of Or85b OSNs in ab3 sensilla for the same strains as in panel **A**. Right: quantifications of Or85b OSNs for the same strains as in panel **A**. **C.** Left: example images of Or10a and Or42b OSNs labelled in ab1 sensilla for the same strains as in panel **A**. Right: quantifications of Or10a and Or42b OSNs for the same strains as panels **A**. For **A**-**C**, letters above boxplots indicate pairwise significant differences (all p < 0.05, Wilcoxon test followed by Bonferroni correction).10 antennae were imaged and quantified for each sensillum type per strain.

In addition to these OSN population expansions, several sensilla and/or OSN types are reduced in *D. sechellia*. For example, ab1 sensilla, which house Or10a and Or42b expressing OSNs, display a nearly 2-fold reduction in number relative to *D. melanogaster*^20,29^. In *D. melanogaster*, Or42b neurons are required for attraction to apple cider vinegar^23^, so their decrease in *D. sechellia* plausibly reflects a lower significance of this sensory stimulus. Labelling of Or10a and Or42b neurons revealed a consistent reduction in all tested *D. sechellia* strains compared to *D. melanogaster* (Figure 3C).

### Adult noni toxin resistance

As part of its specialization on noni, *D. sechellia* has evolved a high resistance to octanoic acid, an abundant noni chemical^30^ that accounts for this fruit’s toxicity for other drosophilids^31,32^. The mechanistic basis of adult octanoic acid resistance in *D. sechellia* is still poorly-understood, but genome-wide association and gene expression studies identify a complex genetic architecture^33^ highlighting candidates involved in cuticle formation^34–36^, enzymatic detoxification^37–39^ and metabolism^40,41^.

To compare the resistance to octanoic acid of our laboratory-adapted strains with that of recent collections, we used a plate-based assay^33^. Adult males and females were combined with 2 μl of pure octanoic acid in a Petri dish to ensure exposure to its volatiles and scored every 5 min for loss of consciousness over 60 min (Figure 4A). When we calculate the cumulative survival probability across 10 replicates for our *D. sechellia* and *D. melanogaster* strains, we observe a consistent species difference: for all *D. sechellia* strains, more than 60% of females (Figure 4B) and males (Figure S1A) remained conscious, while only ∼20% of *D. melanogaster* females and no males did. Comparing the proportion of surviving flies at 60 min between strains for both females (Figure 4C) and males (Figure S1B), all *D. sechellia* strains were significantly different from *D. melanogaster*. For females, we detected some variation among our wild-caught *D. sechellia* strains (Figure 4C), but this reflected slightly lower octanoic acid resistance levels compared to the laboratory strains.

**Figure 4.**
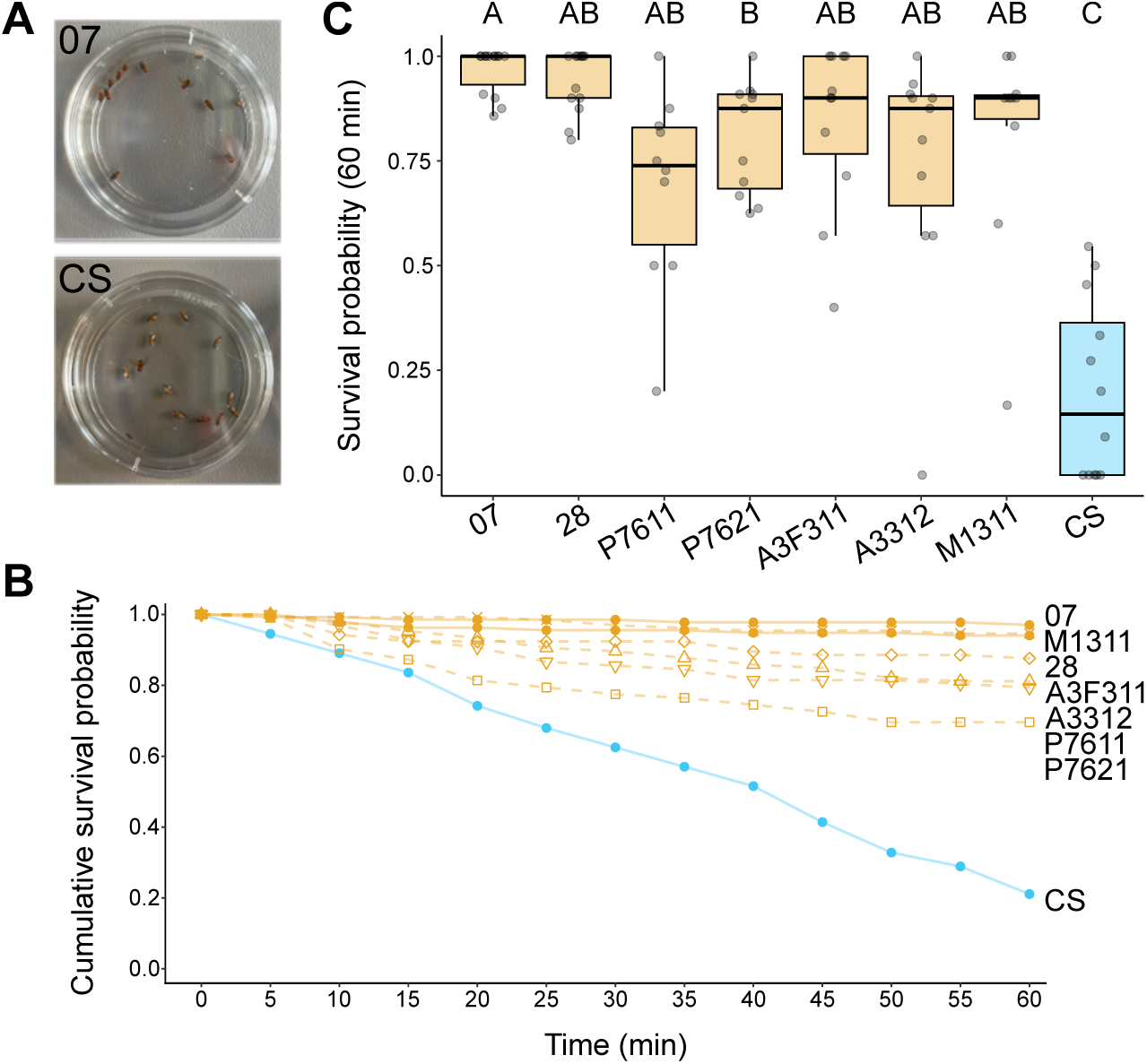
*D. sechellia* adult females display strong resistance to the noni toxin octanoic acid. **A.** Photographs of a single replicate of laboratory *D. sechellia* (07) and *D. melanogaster* (CS) strains following 60 min of exposure to pure octanoic acid. **B.** Cumulative survival probabilities (mean proportion of living flies across all replicates) of laboratory *D. sechellia*, wild *D. sechellia*, and *D. melanogaster* CS is shown for 5 min increments over a 60-min exposure duration. **C.** Final survival probability (proportion of living flies after 60 min of exposure to octanoic acid) for each replicate is shown for wild and laboratory *D. sechellia* strains and *D. melanogaster*. Letters above boxplots indicate pairwise significant differences (all p < 0.05, Wilcoxon test followed by Bonferroni correction). Sample sizes: 07 (14), 28 (13), P7611 (10), P7621 (11), A3F311 (11), A3312 (11), M1311 (10), and CS (12).

### Fecundity and egg size

In addition to adult resistance to noni toxins, *D. sechellia*’s eggs must be able to develop on this toxic fruit. The eggs of *D. sechellia* are larger (∼50% volume) than those of *D. melanogaster*^42^ and it has been hypothesized that this phenotype represents part of the mechanism permitting their successful development on noni. This potential increase in resource allocation toward individual eggs is further hypothesized by some to have resulted in a trade-off with overall fecundity^32,33^, due at least in part to a decrease in total ovariole number^43^, though it remains to be determined if and how this trait evolved during *D. sechellia*’s specialization. While the molecular basis of reduced ovariole number is unknown, association mapping studies have indicated a complex genetic architecture^44,45^, and developmental studies have identified differences in somatic gonadal precursor cell number^46^ and insulin signaling^47^ that contribute to this trait.

To examine these phenotypes in wild-caught flies, we quantified reproductive output for mated females, ovariole number and egg size. We used the single-fly oviposition assay described above to determine the total reproductive output of a single female over a 24-h period. Comparing the total eggs produced per female, irrespective of oviposition media, we found that all *D. sechellia* strains were consistently distinguishable from *D. melanogaster,* laying ∼5-fold fewer eggs (all p < 0.001). Similarly, all *D. sechellia* strains, irrespective of collection date, had fewer ovarioles (∼2-fold reduction) than our *D. melanogaster* strain (Figure 5B). Finally, all *D. sechellia* strains laid larger eggs than *D. melanogaster* (Figure 5C), with the average volume of *D. sechellia*’s eggs being 45% greater than those of *D. melanogaster* Canton-S. For these latter two phenotypes, we did not observe any significant differences between any *D. sechellia* strain (Figure 5B-C).

**Figure 5.**
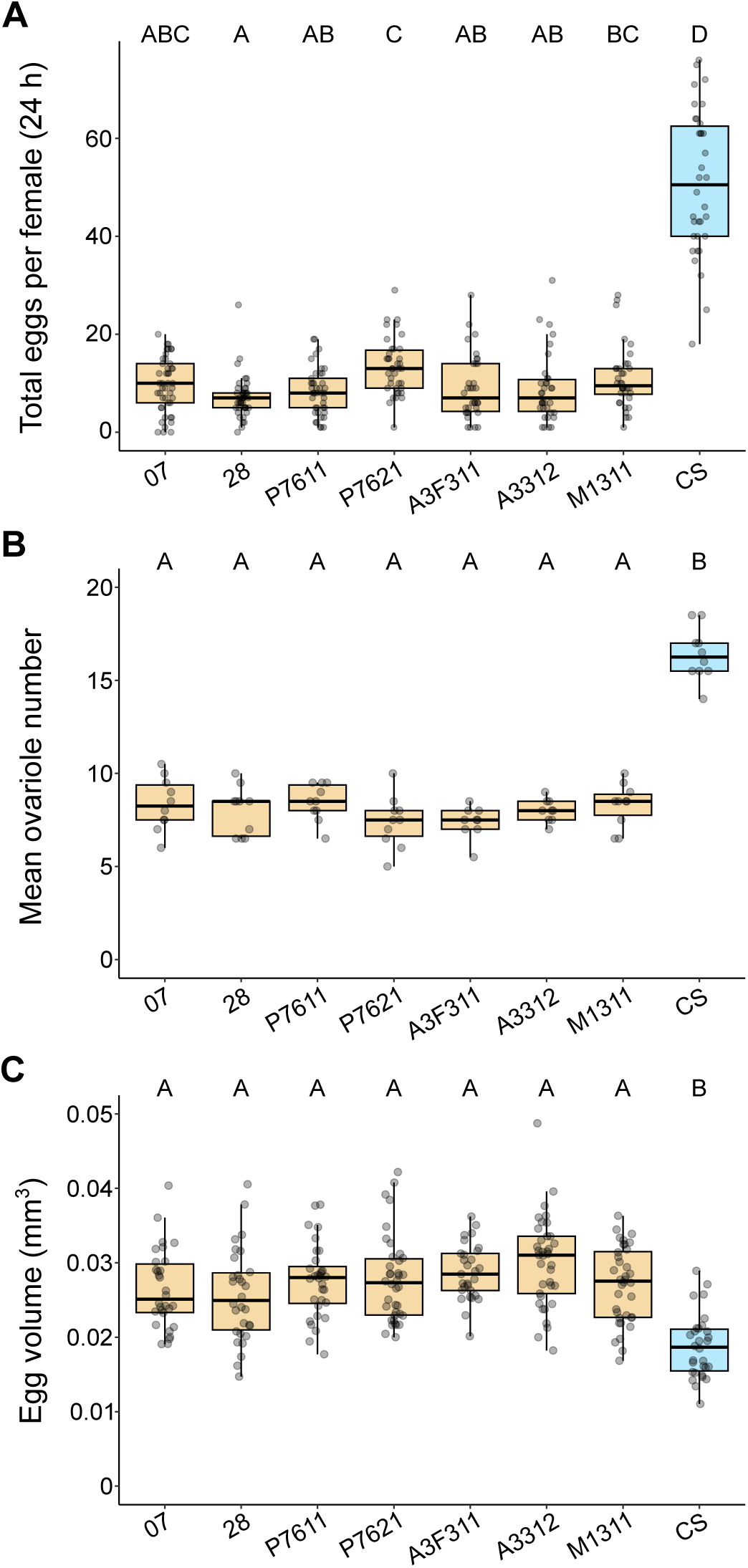
*D. sechellia* lay fewer, larger eggs, and display a reduced ovariole number compared to *D. melanogaster*. **A.** The total eggs laid per female in a 24-h period is shown for laboratory *D. sechellia*, wild *D. sechellia*, and *D. melanogaster* CS. Total eggs laid per female was obtained from the experiments shown in panels 2C-D. **B.** The volume of individual eggs for the same strains as panel **A**. Sample sizes: 07 (30), 28 (30), P7611 (32), P7621 (38), A3F311 (29), A3312 (37), M1311 (36), and CS (30). **C.** The mean number of ovarioles per ovary for individual adult female flies is shown for the same strains as panels **A** and **B**. Sample sizes: 07 (10), 28 (10), P7611 (10), P7621 (10), A3F311 (9), A3312 (9), M1311 (10), and CS (10). For **A**-**C**, letters above boxplots indicate pairwise significant differences (all p < 0.05, Wilcoxon test followed by Bonferroni correction).

Taken together, the decreased fecundity, decreased ovariole number and increased egg size we document is similar to previously described reproductive divergence between laboratory strains of *D. melanogaster* and *D. sechellia*^42,44,48^, and consistent with the hypothesis of an evolutionary trade-off between reproductive output and resource allocation^26^.

### Reduced circadian plasticity

Finally, in addition to *D. sechellia*’s many adaptations to its host fruit, it was recently discovered to be specialized to aspects of its equatorial environment, notably photoperiod (day-length)^49^. Drosophilid flies typically display a bimodal pattern of circadian activity (with some exceptions)^50^, where individual flies time their peak morning and evening activity with sunrise and sunset. Flies living at high latitudes display an ability to plasticly adjust the timing of these peaks to match seasonal variation in day length, with the degree of plasticity correlating with population latitude^49,51,52^. *D. sechellia* are found exclusively at the equator, where day length varies minimally throughout the year^8^. As such, these flies are far less able to delay the timing of their evening peak activity to match longer days that would be experienced at higher latitudes^49^. This loss of plasticity has a polygenic basis, including divergent regulation of the circadian neuropeptide, *Pigment dispersing factor*^49^.

We used the *Drosophila* activity monitor system to compare the activity patterns of our strains under both equatorial and high latitude summer conditions^53^. We first recorded male flies of each strain for 7 days under 12:12 h LD (equatorial) conditions at 25°C (Figure 6A). All strains of both species displayed a characteristic peak of activity surrounding subjective dawn and dusk, separated by an afternoon period of reduced activity. We next took the average activity for each fly per strain during the last 4 days of this recording and annotated the timing of the evening activity peak (Figure 6B). *D. melanogaster* and all the laboratory and wild-caught *D. sechellia* strains displayed a high synchronicity of their evening peak time with the timing of subjective dusk (12 hours).

**Figure 6.**
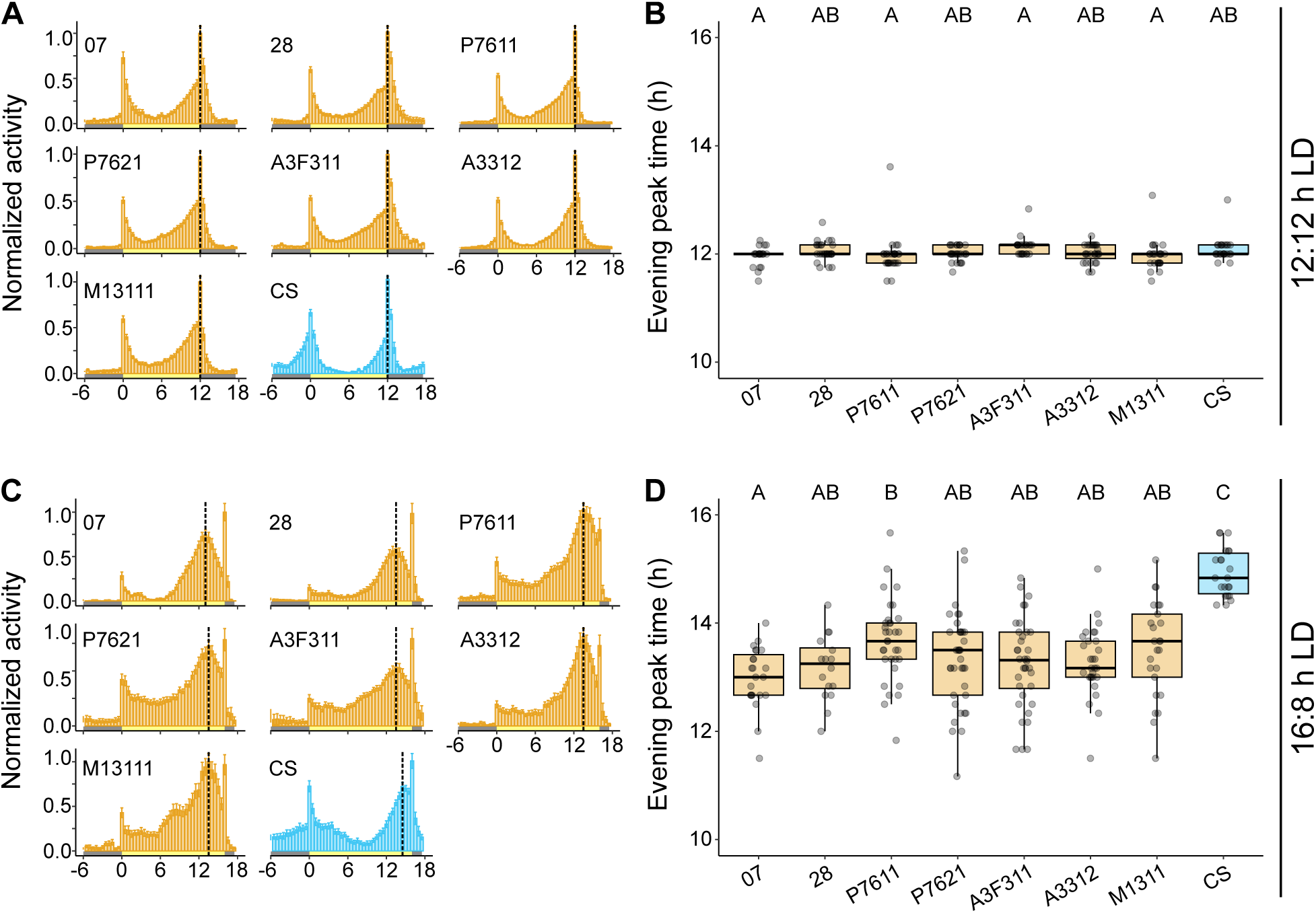
*D. sechellia* displays reduced circadian plasticity during long days. **A.** Maximum normalized average activity plots for laboratory *D. sechellia*, wild *D. sechellia*, and *D. melanogaster* CS under a 12:12 h photoperiod. **B.** Mean evening peak time for the last 4 days of a 7-day period under 12:12 h photoperiod conditions is shown for individual flies for the same strains as panel **A**. Sample sizes for panels **A** and **B**: 07 (21), 28 (28), P7611 (29), P7621 (28), A3F311 (24), A3312 (27), M1311 (26), and CS (24). **C.** Maximum normalized average activity plots for the same strains as panels **A** and **B** under a 12:12 h photoperiod. **D.** Mean evening peak time for the last 4 days of a 7-day period under 16:8 h photoperiod conditions is shown for individual flies for the same strains as panels **A**-**C**. Sample sizes for panels **C** and **D**: 07 (23), 28 (16), P7611 (34), P7621 (37), A3F311 (36), A3312 (29), M1311 (25), and CS (22). For **A** and **C**, histograms display maximum normalized activity of flies in 30-min bins across a 24-h period. Error bars denote standard error of the mean. Grey and yellow bars along the x-axis represent lights-off and lights-on in the incubator, respectively. Vertical dashed lines represent the average evening peak time. For **B** and **D**, letters above boxplots indicate pairwise significant differences (all p < 0.05, Wilcoxon test followed by Bonferroni correction).

We next modified the photoperiod for these flies, recording them for an additional 7 days under 16:8 h LD (high latitude summer) conditions (Figure 6C). As before, we used the average activity from the last 4 days of this recording period from each fly to determine the timing of the evening activity peak under extended photoperiod conditions (Figure 6D). As previously described^49^, we observed a clear species difference in evening peak plasticity, where our *D. melanogaster* laboratory strain was able to delay its evening peak time significantly longer than any of our *D. sechellia* strains. On average, *D. melanogaster* Canton-S was able to delay its evening peak to ∼15 h, just 1 h shy of subjective dusk. By contrast, our *D. sechellia* strains’ average evening peak time ranged from ∼13-13.67 h. Comparing laboratory and wild-caught *D. sechellia* strains identifies a single wild-caught strain from Praslin Island (P7611) with a slightly elevated evening peak time relative to just one of our laboratory strains, *Dsec*07. There is therefore no consistent discernable difference in circadian plasticity between laboratory and wild-caught *D. sechellia* strains. This is unsurprising because, for this phenotype, standard laboratory fly culture conditions (12:12 h LD cycle at 25°C) more accurately reflect equatorial conditions than the variable conditions at high latitudes in the wild.

## Discussion

Here, we sought to detect potential effects of laboratory adaptation on several phenotypes of *D. sechellia*. Our results indicate that *D. sechellia* laboratory strains do not display signs of a loss or degradation of the behavioral or anatomical phenotypes of this species, whose evolution was likely influenced by its specialization to a specific ecological niche. We discuss these results on a phenotype basis below.

Despite not being reared in the presence of noni fruit for hundreds of generations, *D. sechellia* laboratory strains show a strong retention of preference for noni-based substrates in both olfactory and oviposition assays, along with the increases/decreases in OSN number associated with these preferences. For this study, we maintained our *D. sechellia* strains on standard *Drosophila* media supplemented with a paste made with commercially available noni juice. It is possible that this approximation of noni fruit is sufficient to maintain noni preferences during laboratory culture. However chemical comparison of this juice with ripe noni fruit reveals differences in the relative proportions of specific chemicals, such as octanoic and hexanoic acids^14^. Our laboratory *D. sechellia* strains were originally sourced from the *Drosophila* Species Stock Center (DSSC), where they were previously maintained for years on a cornmeal media (DSSC Director Patrick M. O’Grady, *personal communication*). Given the many known examples of laboratory adaptation, that phenotypes can be experimentally evolved in far fewer generations^40,54,55^, and the numerous examples demonstrating that such preferences are highly evolvable^14,21,26^, why, then, have the *D. sechellia* laboratory strains studied here not evolved novel preferences to laboratory media? It is possible that these isogenic strains have simply not been reared in the laboratory for long enough to evolve altered preferences de novo. Alternatively, a laboratory environment might not select strongly for these preferences: in standard culture vials, flies have no need to forage and are not given a choice between media, and therefore have little option to exert (or need for) a preference. Thus, any chemosensory preferences might be effectively neutral in a laboratory setting, and such phenotypes would only degrade slowly over time via the accumulation of random mutations and genetic drift.

This hypothesis might also explain the persistence of octanoic acid resistance in laboratory populations. There is no obvious benefit to retaining octanoic acid resistance in the lab, as these flies never encounter it. However, as long as the cost of retaining octanoic acid resistance is not large, selection also would not favor its loss, rendering it a neutral phenotype. Alternatively, there might be hidden selective advantages. For example, cuticle changes that confer octanoic acid resistance might also result in decreased desiccation risk, which could still be favored in a laboratory setting. Thus, laboratory selection on one effect of the phenotype might maintain the other.

We also show that the reduced fecundity and ovariole number previously described in *D. sechellia* laboratory strains is indeed representative of the phenotypes of wild-caught flies. Laboratory environments typically select for high fecundity through high-density rearing practices^3^. Our laboratory strains do not appear to have responded to this selection pressure. However, if the hypothesis of a trade-off in *D. sechellia* of reduced fecundity with increased egg size^32,33^ is correct, for laboratory strains to evolve increased fecundity, they might also have to evolve reduced egg size, which presents a more complicated evolutionary trajectory. In this respect, reduced fecundity of *D. sechellia* might represent a evolutionary dead-end. Future studies exploring differences in maternal investment to directly test this hypothesis seem warranted.

Laboratory *D. sechellia* strains have also retained their circadian specialization to an equatorial environment. Considering most laboratories maintain flies under equatorial conditions with respect to photoperiod and temperature, this is unsurprising. In this context, it is notable *D. melanogaster* maintained at equatorial conditions have not lost their capacity for photoperiodism, generally supporting the idea that laboratory strains maintain, at least over these time scales, aspects of local adaptation to their collection site.

Finally, while it is most parsimonious to assume that the mechanisms underlying species-specific traits are conserved between laboratory and wild-caught strains – as supported here by the described expansions and contractions of OSN numbers – future studies should further investigate in our wild-caught strains the previously-described evolution of gene expression^37,38,40,49,56,57^, the nervous system^14,20,21,24,26,49,57,58^, and the reproductive system^46,47^. Regardless, our data demonstrate robustness in *D. sechellia*’s specialism-associated phenotypes, further supporting the power of this species as a model system to uncover ecologically relevant genetic adaptations.

## Methods

### *D. sechellia* wild-caught strain maintenance

Following field collections in April 2023, isofemale *D. sechellia* lines were established and maintained in the Extavour laboratory on *Drosophila* media (0.8% agar, 2.75% yeast, 5.2% cornmeal, 11% dextrose) without noni juice at 25°C, 65% relative humidity, and a 12:12 h light-dark cycle, from May-November 2023. From November 2023 onward, flies from each strain were reared in the Benton laboratory on *Drosophila* media (0.6% agar, 5.8% yeast, 5.8% wheat, 10% fruit juice) supplemented with noni juice paste (Raab Vitalfood Bio noni juice combined with instant *Drosophila* medium, Carolina #173210). From January 2025 onward, flies were reared in the Shahandeh laboratory on *Drosophila* media (0.9% agar, 2.6% yeast, 6.25% cornmeal, 6.25% molasses) supplemented with noni juice (Healing Noni juice combined with instant *Drosophila* medium, Carolina #173210). In both the Benton and Shahandeh laboratories, flies were maintained in non-overlapping 2-week generations at 25°C, 50% relative humidity, and a 12:12 h light-dark cycle. To limit the potential for laboratory adaptation to occur, all behavioral analyses reported herein were conducted within a year of collection (by May 2024) in the Benton laboratory. Quantifications of ovariole number, egg size (Shahandeh laboratory), and OSN number (Benton laboratory) were conducted from July-December 2025.

### *Drosophila* laboratory strain maintenance

The laboratory strain of *D. melanogaster* (Canton-S, RRID:BDSC_64349) was maintained on a *Drosophila* media (recipes above) in the Benton and Shahandeh laboratories in non-overlapping 2-week generations at 25°C, 50% relative humidity, and a 12:12 h light-dark cycle. The laboratory strains of *D. sechellia* (*Dsec.*07, DSSC 140210248.07; *Dsec.*28, DSSC 140210248.28) were maintained under the same conditions on *Drosophila* media supplemented with noni juice paste (as described above for each laboratory).

### Species identification by molecular barcoding

Wild-caught flies were identified as *D. sechellia* using molecular barcoding of portions of the *mitochondrial cytochrome c oxidase* (*mt:CoI*) gene, using COI-218F (5’ CAACATTTATTTTGATTTTTTGG 3’) and COI-3037R (5’ TYCATTGCACTAATCTGCCATATTAG 3’) primers, using standard methods. In brief, genomic DNA was extracted from individual flies, amplified via PCR using OneTaq 2X Master Mix (includes 0.62 units Taq; New England BioLabs #M0480), and purified using Monarch PCR & DNA Cleanup Kit (New England BioLabs #T1030L). The resulting products were subject to Sanger sequencing (Bioscience Inc), using COI-218F primer. Raw sequence data were analyzed using Chromas Lite v2.6.6 (Technelysium Pty Ltd. Australia) and sequences were compared with those in GenBank to identify the species. Flies were identified as *D. sechellia* if the *mt:CoI* amplicon showed at least 99.5% sequence identity with GenBank accession MK659840.1, which is the *mt:CoI* sequence of *D. sechellia* voucher DSECH20161109 complete mitochondrial genome. *mt:CoI* amplicon sequences from the strains described in this study were deposited in GenBank (Accession numbers: M1311, PX502215; A3312, PX502212; A3F311, PX502211; P7611, PX502214; P7621, PX502213).

### Olfactory trap assay

The olfactory trap assay was replicated from previous studies investigating *D. sechellia*’s short-range olfactory preference^14,21^. In brief, 25 3-5 day-old, mated females were anesthetized with ice and added to a 12 cm circular plastic Tupperware containing two olfactory traps prior to sealing the tops with a fine mesh secured by rubber band (Figure 2A). Each trap contained 300 μl of either noni juice (Raab Vitalfood) or apple cider vinegar (Denner brand) mixed with Triton-X (final concentration 0.2%). Flies can enter the traps through a pipette tip puncturing the cap but are unable to find the opening to escape after selecting an odor source. Traps were stored for 24 h at 25°C and ∼60% humidity under red light, after which the number of flies in each trap, or remaining untrapped animals (live and dead), were counted. We excluded from our statistical analyses any traps where more than 25% of flies remained untrapped (Figure S2). We calculated preference indices as: (number of flies in the noni juice trap − number of flies in the apple cider vinegar trap)/number of living flies, both trapped and untrapped. We used a Wilcoxon rank-sum test to test individual preference indices for differences from 0 (no preference), and pairwise Wilcoxon tests to compare preference indices between strains. For both, p-values were corrected for multiple comparisons using the Bonferroni method.

### Two-choice single-fly oviposition assay

To measure individual oviposition preferences, we used a single-fly chamber assay where 30 3-5 day old mated females are housed individually in plexiglass cells where they can sample two oviposition media^24,27^ (Figure 2C). We provided a choice between two substrates: a 0.67% agar solution made with noni juice, or an agar solution of the same density prepared with apple cider vinegar. Prior to assay, females were housed in high densities with males for 3 days to ensure they were gravid. The chambers were housed at 25°C and ∼60% humidity for 24 h under a 12:12 h light-dark cycle. At the end of the assay, flies were removed and the number of eggs present on each substrate was recorded for each chamber. We calculated preference indices as: (number of eggs laid on the noni media – the number of eggs laid on apple cider vinegar)/total number of eggs laid. We used a Wilcoxon rank-sum test to test individual preference indices for differences from 0 (no preference), and pairwise Wilcoxon tests to compare preference indices between strains. For both, p-values were corrected for multiple comparisons using the Bonferroni method.

### Histology and imaging

Hybridization chain reaction RNA fluorescence in situ hybridization (HCR-FISH) was performed as described in^59^. RNA probes for *Or10a* (accession number NM_078567.5), *Or42b* (NM_078900.2) and *Or85b* (NM_079555.2), amplifiers, and buffers were purchased from Molecular Instruments (HCR^TM^ RNA FISH (v3.0)). Probe sequences correspond to *D. melanogaster Or* genes, but have ≥95% identity with orthologous *Or*s in *D. sechellia*.

Immunofluorescence for Ir75b was performed as described previously (Auer et al. 2020), using anti-Ir75b (Prieto-Godino et al., 2016) diluted 1:500. Secondary anti-guinea pig antibody conjugated to Alexa Fluor 488 (A11073, Invitrogen AG) was diluted 1:1000.

Images were acquired using a Zeiss confocal microscope LSM 710 with a 40× objective and processed in ImageJ. Neurons were counted manually using the ImageJ MultiPoint tool.

### OA resistance assay

To measure adult resistance to octanoic acid, we used a previously-described plate assay^33,40^ (Figure 3A). In brief, ten male or female flies were briefly anesthetized with CO_2_ and placed in a 60 mm diameter × 15 mm high Petri dish to which 2 μl of pure octanoic acid had been previously added to the center of the lid. Upon recovery from anesthesia, the flies were scored every 5 min for loss of consciousness for 60 min. To test for differences between strains, we compared the proportion of surviving flies at the final time point using pairwise Wilcoxon tests followed by post-hoc correction for multiple comparisons.

### Ovary dissection and counting

3-5-day-old, mated females (reared at 25°C, ∼60% relative humidity, 12:12 h LD) were anesthetized on CO_2_ and sorted by sex. Whole flies were fixed for 2 h at room temperature in 4% paraformaldehyde in PBS containing 0.3% Triton X-100 (PBS-TX; 1 ml per ∼30 flies) on a nutator. Samples were washed 3 × 5 min in PBS-TX and ovaries were dissected in PBS-TX under a stereomicroscope. To enhance contrast for ovariole visualization, each set of dissected ovaries was briefly dipped in crystal violet solution (K1184, APExBio). Each ovary was gently teased apart in PBS-TX on a ceramic slide using an eyelash probe to separate ovarioles without rupturing the ovary. Ovarioles were identified as individual strings of progressively developed egg chambers. Both ovaries from each female were counted and averaged to obtain mean ovarioles per ovary.

### Egg collection and size measurements

Oviposition plates were prepared as 5% sucrose, 2% agarose in distilled water and diluted with noni juice or apple cider vinegar to a final concentration of 1%. Plates were poured into 60 mm diameter Petri dishes and cooled to room temperature. Groups of mated females (3-5 days old; conditioned with males at high density for 3 days prior) were transferred to fresh plates and allowed to oviposit for ∼20 h at 25°C, ∼60% relative humidity, 12:12 h light-dark cycle. For imaging, a dissecting scope was used to locate eggs; individual eggs were placed into a single file on the plate on their dorsal side. We photographed each plate along with a ruler for scale using a Nikon Z50 DX 16-50mm on a tripod approximately 24 cm above the plate’s surface. Egg size was quantified in Fiji/ImageJ (https://imagej.net/ij/). For each egg, length and width were measured using the straight-line tool. We calculated total egg volume using a previously established formula^60^: V = 1/6πW^2^L for ∼30 eggs of each strain. Two observers independently estimated egg volume to control for observer bias (Figure 5C and Figure S3), as a completely blind study was not possible due to differences in the color of substrate used for each species.

### *Drosophila* activity monitoring

We used the *Drosophila* Activity Monitor system to record the daily activity patterns of our strains under both equatorial (12:12 h LD) and high latitude summer (16:8 h LD) photoperiod conditions^61^, replicating experiments used to describe reduced circadian plasticity in *D. sechellia*^49^. Briefly, we exposed individual 3-5 day old males first to 7 days of an equatorial photoperiod, followed by 7 days of a high latitude summer photoperiod. We used a previously developed algorithm to quantify the timing of each fly’s evening peak^49^ using the last 4 days of each 7-day period, discarding the first three days as an acclimation period. We compared evening peak time between our strains using pairwise Wilcoxon tests followed by correction for multiple comparisons.

To estimate the circadian period of each strain, we acclimated flies to a 12:12 h LD cycle for 4 days before subjecting them to 5 days of constant darkness conditions. We used the constant darkness behavioral data to calculate the duration of each strain’s free-running period with the R package Rethomics, which was also used for all circadian data visualization^62^. We compared periods across strains with pairwise Wilcoxon tests subjected to post-hoc correction for multiple comparisons.

### Statistical analysis and data visualization

All statistical analyses were conducted using the ‘stats’ package (v4.6.0) in R (v4.4.2)^63^. Data were visualized using ‘ggplot2’^64^. For boxplots in all figures, the bold line represents the median value, with the length of the box displaying the interquartile range. Whiskers signify the last quartiles excluding outliers.

## Acknowledgements

We thank the Seychelles Bureau of Standards, the Seychelles Ministry of Agriculture, Climate Change and Environment, and the Seychelles Islands Foundation for permission to collect and perform research with field-collected samples of *Drosophila* and *Morinda citrifolia* fruits from the Seychelles. We thank Joseph Francois and Gérard Rocamora of the University of Seychelles for invaluable assistance with field collections, and Pat Mathiot for discussions of the evolution and diversity of endemic Seychelles flora and fauna. Research in C.G.E.’s laboratory was supported by funds from Harvard University and the Howard Hughes Medical Institute (HHMI). C.G.E. is an HHMI Investigator. Research in R.B.’s laboratory was supported by the University of Lausanne, an ERC Advanced Grant (833548) and the Swiss National Science Foundation (310030_219185). Research in M.S.’s laboratory was supported by Hofstra University.

---

**Supplemental Figure 1.**
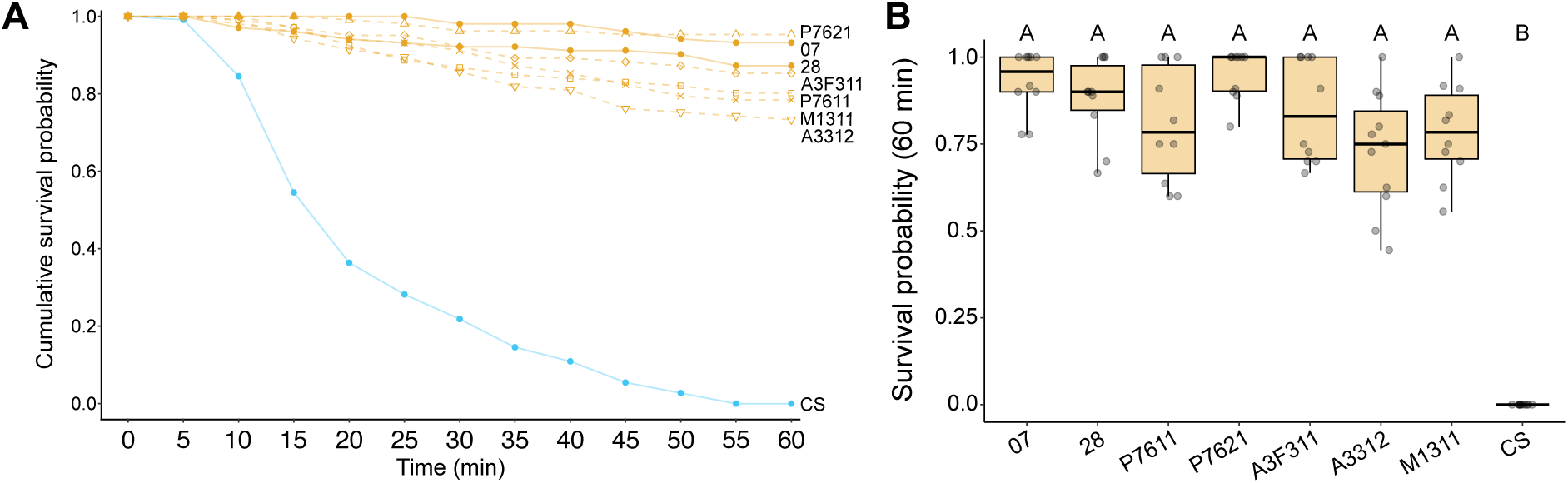
*D. sechellia* adult males display strong resistance to the noni toxin octanoic acid. **A.** Cumulative survival probabilities (mean proportion of living flies across all replicates) of laboratory *D. sechellia*, wild *D. sechellia*, and *D. melanogaster* CS is shown for 5-min increments over a 60-min exposure duration. **B.** Final survival probability (proportion of living flies after 60 min of exposure to octanoic acid, y-axis) for each replicate is shown for wild and laboratory *D. sechellia* strains and *D. melanogaster* (x-axis). Letters above boxplots indicate pairwise significant differences (all p < 0.05, Wilcoxon test followed by Bonferroni correction). Sample sizes: 07 (10), 28 (10), P7611 (10), P7621 (10), A3F311 (10), A3312 (11), M1311 (10), and CS (10).

**Supplemental Figure 2.**
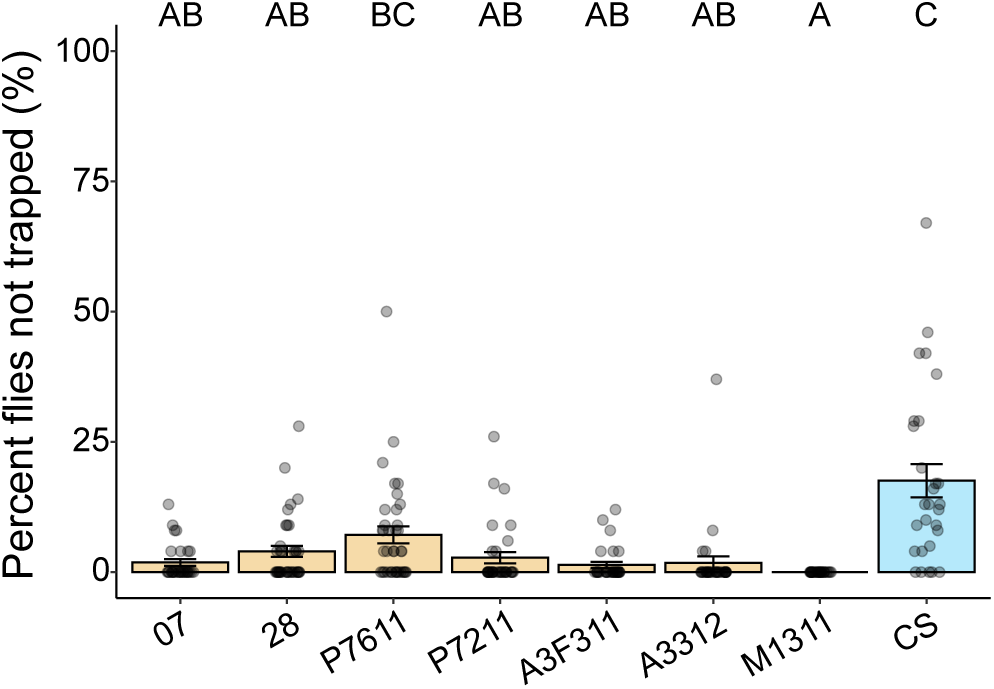
Percent of flies not trapped during short-range olfactory trap assay. The percent of flies that were found in neither the noni juice nor apple cider vinegar trap for each replicate for laboratory *D. sechellia*, wild *D. sechellia*, and *D. melanogaster* CS. Bar plots represent mean values and error bars denote standard error of the mean. Letters above boxplots indicate pairwise significant differences (all p < 0.05, Wilcoxon test followed by Bonferroni correction). Sample sizes are provided in Figure 2 legend.

**Supplemental Figure 3.**
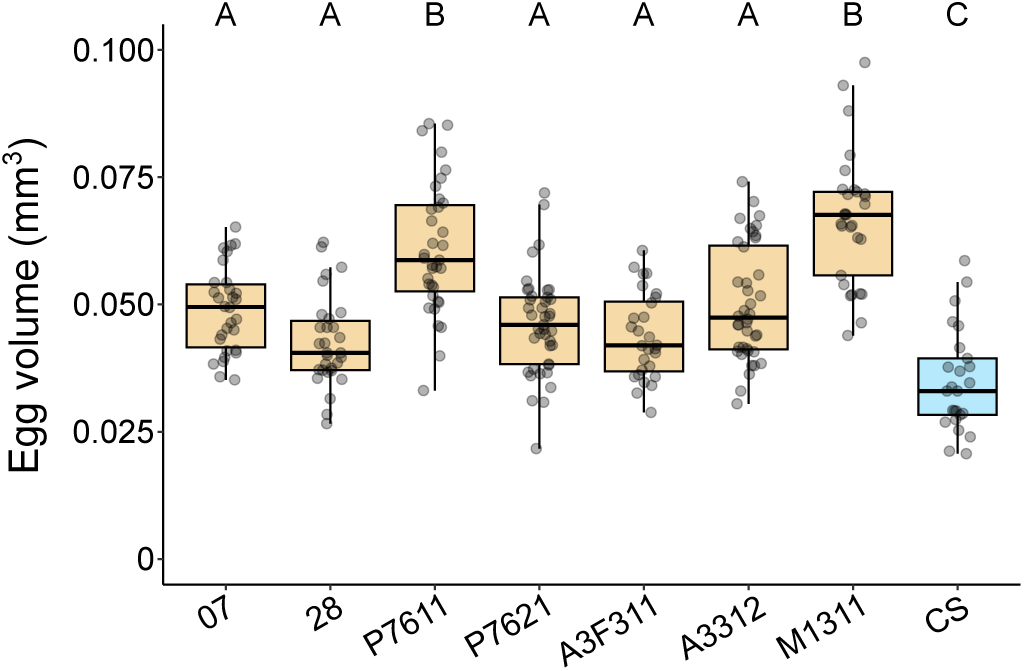
Egg volume quantified by a second observer confirms *D. sechellia* lays larger eggs than *D. melanogaster*. The volume of individual eggs is displayed for laboratory *D. sechellia*, wild *D. sechellia*, and *D. melanogaster* CS. Letters above boxplots indicate pairwise significant differences (all p < 0.05, Wilcoxon test followed by Bonferroni correction). Sample sizes: 07 (30), 28 (30), P7611 (35), P7621 (41), A3F311 (28), A3312 (40), M1311 (30), and CS (25).

